# Pre-clinical testing of two serologically distinct chimpanzee-origin adenovirus vectors expressing spike of SARS-CoV-2

**DOI:** 10.1101/2022.02.23.481620

**Authors:** Mikhail Novikov, Mohadeseh Hasanpourghadi, Robert Ambrose, Arezki Chekaoui, Dakota Newman, Wynetta Giles-Davis, Zhiquan Xiang, Xiang Yang Zhou, Hildegund CJ Ertl

## Abstract

Two serologically distinct chimpanzee-origin, replication-defective adenovirus (AdC) vectors expressing the spike (S) protein of an early severe acute respiratory syndrome coronavirus 2 (SARS-CoV-2) isolate were generated and tested for induction of antibodies in young and aged mice. Both vectors induced S protein-specific antibodies including neutralizing antibodies. Levels of antibodies increased after a boost. The effectiveness of the boost depended on vector dose and timing between the two immunizations. Using two heterologous AdC vectors was more effective than vaccinating with the same vector repeatedly. Antibodies partially crossreacted between different S protein variants. Cross-reactivity increased after booster immunization with vectors carrying the same S gene, expression of two different S proteins by the AdC vectors used for the prime and the boost did not selectively increase responses against the variants.

## INTRODUCTION

The global COVID-19 pandemic caused by SARS-CoV-2 killed millions of people worldwide. After the virus was first identified and sequenced at the beginning of 2020^1,2^ vaccines were developed rapidly and entered clinical trials a few months later and within less than one year after onset of the pandemic several gained emergency approval. Those that became first available in the US or Europe are based on mRNA vaccines^3,4^ or replication-defective adenovirus (Ad) vectors derived from human (HAdV)^5,6^ or chimpanzee^7^ serotypes. Additionally inactivated SARS-CoV-2 vaccines were generated and used in South America, Asia, and Africa.^8,9^ All recombinant COVID-19 vaccines used for mass vaccination thus far express the S protein of an early isolate. The S protein binds to the viral receptor, i.e., angiotensin-converting enzyme 2 (ACE2), and is the target for virus neutralizing antibodies (VNAs),^10^ which were shown to protect against infection.^11^

COVID-19 vaccines based on two doses of mRNA are highly efficacious and protect >90% of vaccine recipients.^11,12^ The efficacy of the Ad vector vaccines varies; the one dose J&J vaccine shows 67% protection against disease,^13^ the chimpanzee-derived Ad (AdC) vaccine from AstraZeneca that uses the same vector twice has an efficacy of 76%^14^ while the two dose Sputnik V vaccine that primes with one human serotype Ad vector and then boosts with a serologically distinct Ad vector is as efficacious as the mRNA vaccines.^8^

Although the SARS-CoV-2 vaccines are highly efficacious, they primarily prevent disease rather than infection and protection declines after a few months.^15–20^ A boost at 6 months after the initial immunization with one of the most commonly used vaccine prototype, i.e., the mRNA vaccines, increases VNA titers albeit only temporarily potentially necessitating another boost as soon as 4 months later.^21^ Although an additional boost was shown to restore protection, antibody titers failed to exceed those after the 3^rd^ boost.^22^ Confounding the problem of short-live immunity after COVID-19 vaccination has been the emergence of more transmissible viral variants that partially escape the vaccine-induced immune responses including those after 4 doses of the mRNA vaccines.^22–25^

Limited pre-clinical studies were conducted with the COVID vaccines to optimize immune responses regarding potency and longevity by testing different vaccine doses, varying timing between the prime and the boost or systematically exploring the use of heterologous vaccine platforms. The short-lived efficacy of the most used COVID-19 vaccines is of concern; it is not feasible to vaccinate the human population every 4 to 6 months. Additional studies to develop second generation vaccines or vaccine regimens, which induce more sustained protective immunity are needed.

Here we describe pre-clinical results with two serologically distinct AdC vectors expressing the S protein of an early SARS-CoV-2 isolate. Our results show that the vaccines induce S protein-specific antibodies including VNAs in young and aged mice. Antibody responses increase after a boost and are then maintained at stable levels for at least 5 months. The magnitude of the booster response depends on vaccine dose and interval between the two immunizations and is enhanced by using heterologous vectors. VNAs generated by a single vaccine dose show limited responses to the S protein of viral variants; cross-reactivity increases after a boost. Vaccinating mice sequentially with two AdC vectors expressing either the S protein of the early SARS-CoV-2 isolate, or a version in which the receptor binding domain (RBD) had been mutated to resemble the South African variant followed by a boost with an AdC vector expressing the early S protein only marginally improves cross-reactivity of VNAs to additional viral variants.

## RESULTS

### Generation and testing of AdC-S vaccine vectors

We developed E1-deleted AdC6 (SAdV-23) and AdC7 (SAdV-24) vectors expressing the S protein of an early SARS-CoV-2 isolate from Sweden called SARS-CoV-2/human/SWE/01/2020 (AdC-S_SWE_, GenBank number: QIC53204). In addition, we generated an AdC6 vector expressing the same S protein in which the RBD sequence was modified to incorporate the E484K and N501Y mutations of the B1.351 South African variant (AdC6-S_SWE/B1.351_). Vectors were produced with the help of available molecular Ad clones^26^ and upon expansion, purification, titration and passing quality control assays they were tested for protein expression upon transfection of HEK-293 cells. As shown in Suppl. Fig. 1A both types of AdC vectors express a protein of the expected size that binds the anti-S protein antibody.

### Generation and testing of vesicular stomatitis virus (VSV) vectors pseudotyped with the S protein of SARS-CoV-2

To allow for testing of VNAs we developed several green fluorescent protein (GFP)-expressing VSV vectors that were pseudotyped with S protein of SARS-CoV2 (VSV-S).^27^ One carries the same S protein as the vaccine vectors (VSV-S_SWE_). In a second the S_SWE_ sequence was modified to incorporate the E484K and N501Y RBD mutations of the B1.351 South African variant (VSV-S_SWE/B1.351_). Others express either the S protein with N501Y and P681H RBD mutations present in the B1.1.7 UK variant (VSV-S_SWE/B1.1.7_), or L452R and D614G mutations of the Indian B.1.617.2 delta variant (VSV-S _SWE/B.1.617.2_).

To establish a neutralization assay with the VSV-S vectors, we tested 3 different cell lines including the commonly used VeroE6 cell line but decided on baby hamster kidney (BHK)-21/WI-2 cells, as they gave more consistent results. BHK-21/WI-2 cells express the ACE2 receptors that is used by SARS-CoV2 for cell entry as was shown by immunofluorescent staining and flow cytometry (Suppl. Fig. 1B). VSV-S_SWE_ virus transfection of BHK-21/WI-2 cells is prevented by human antibodies from individuals, who experienced a SARS-CoV-2 infection but not by antibodies from sera of noninfected human controls (Suppl. Fig. 1C). It is also neutralized by sera from S protein-immune but not naïve mice (Suppl. Fig. 1D).

### Antibody responses to different doses of AdC-S vectors

Groups of outbread ICR mice (5/group) were immunized with 10^9^, 10^10^, or 5 x 10^10^ virus particles (vp) of the AdC6-S_SWE_ and AdC7-S_SWE_ vectors. Mice that received the lowest vector dose were boosted 4 weeks after the prime; the other mice were boosted 6 weeks after the prime. All mice were boosted with heterologous AdC vectors given at the same doses that had been used for priming. Mice were bled 2 and or 4 weeks after the prime and 2 weeks after the boost. Mice that received the 1 x 10^10^ vp doses were kept for an additional 5 months and bled at 3 and 5 months after the boost (19 and 27 weeks after the prime) to determine duration of antibody responses (Fig. 1A). Sera from individual mice were tested by an enzyme-linked immunosorbance assay (ELISA) on a mixture of commercially available S1 and S2 proteins of SARS-CoV-2. By 2 weeks after the prime mice in the high dose group developed detectable S protein-specific antibodies while by 4 weeks all animals but for one mouse in the low dose AdC6-S_SWE_ group seroconverted (Fig.1B). Response magnitude was dose dependant. Responses significantly increased by 2 weeks after the boost in most groups (Fig 1C). Antibodies as tested for the 10^10^ vp vaccine groups were maintained at stable levels for at least 5 months after the boost. Antibody isotypes were tested using pooled sera from different time points at a 1:100 dilution. Responses were dominated by antibodies of IgG2 isotypes indicative of the T helper cell type 1 response (Fig. 1D).

**Fig. 1.**
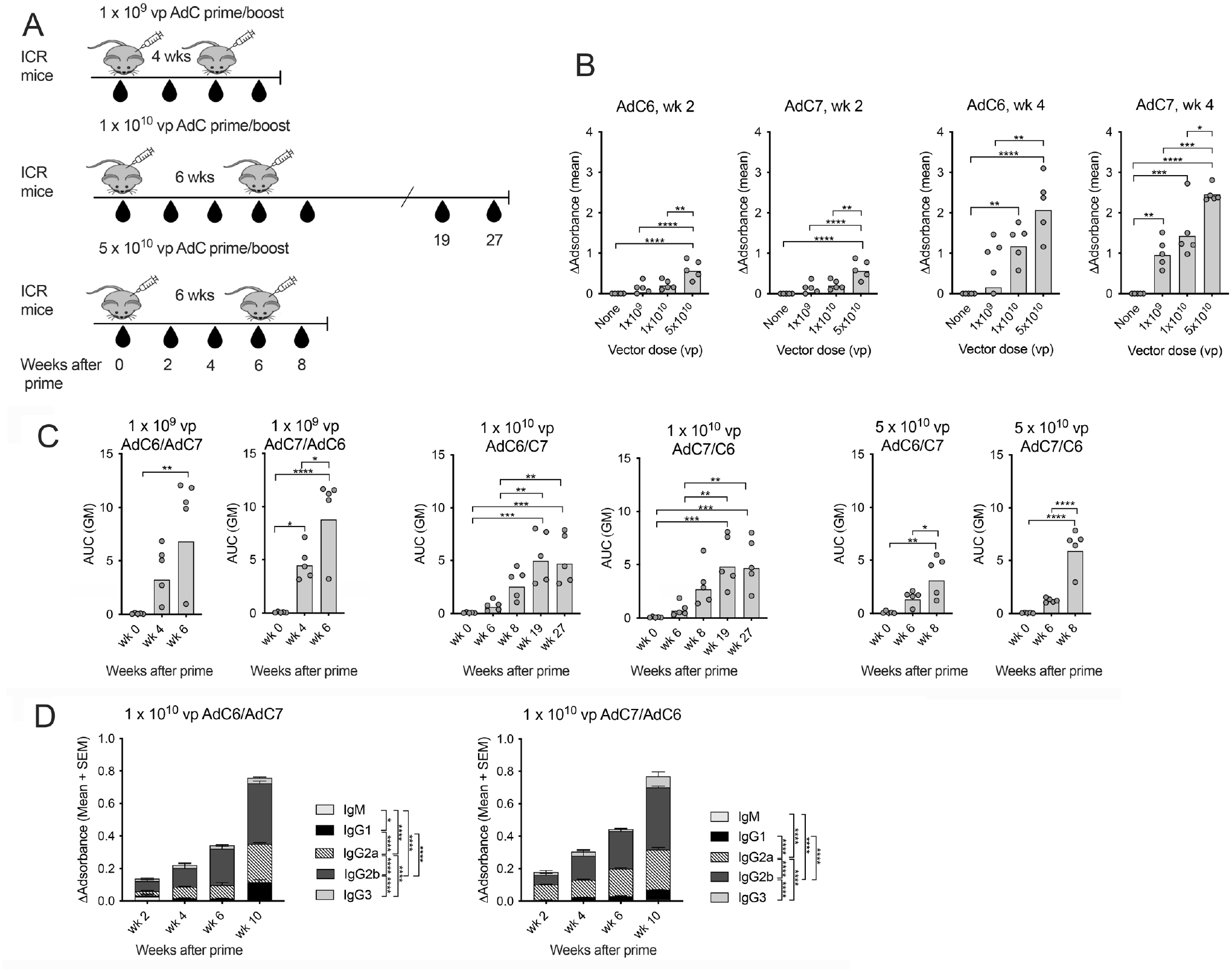
Antibody responses to different doses of the AdC-S vectors. [A] Experimental outline. The drops indicate bleeds. [B] Optical density reading for the ELISA testing sera at a 1:100 dilution from individual mice at 2 and 4 weeks after the initial immunization. Background data without sera were subtracted. Bars indicate geometric means (GM). Data were analyzed by One-way Anova with Tukey’s multiple correction. In this and subsequent graphs lines with stars above indicate significant differences. (*) P value between 0.01-0.05, (**) p-value between 0.001-0.01, (***) p-value between 0.0001-0.001, (****) p-value < 0.0001. [C] Antibody titers as shown as area under the curve of the background corrected OD reading of serially diluted sera. Differences between groups were analyzed by one-way Anova [D] Antibody isotypes of sera harvested at different times from mice primed with 10^10^ vp of the AdC6-S_SWE_ or the AdC7-S_SWE_ vectors. For the image corrected OD values (after subtraction of background data) are stacked. Significant differences calculated by 2-way ANOVA with Tukey’s correction are shown at the side next to the legends.

Some sera from the same mice were tested for neutralization of SARS-CoV-2 virus (Figure 2A). Sera from mice immunized with 1 × 10^10^ or 5 × 10^10^ vp of the AdC vectors were analyzed by a surrogate neutralization assay that tests sera for inhibition of binding of human ACE2 to S protein’s RBD^28^ (Fig. 2B). All mice developed RBD-specific antibodies by week 10, which were maintained without significant declines through week 27.

**Fig. 2.**
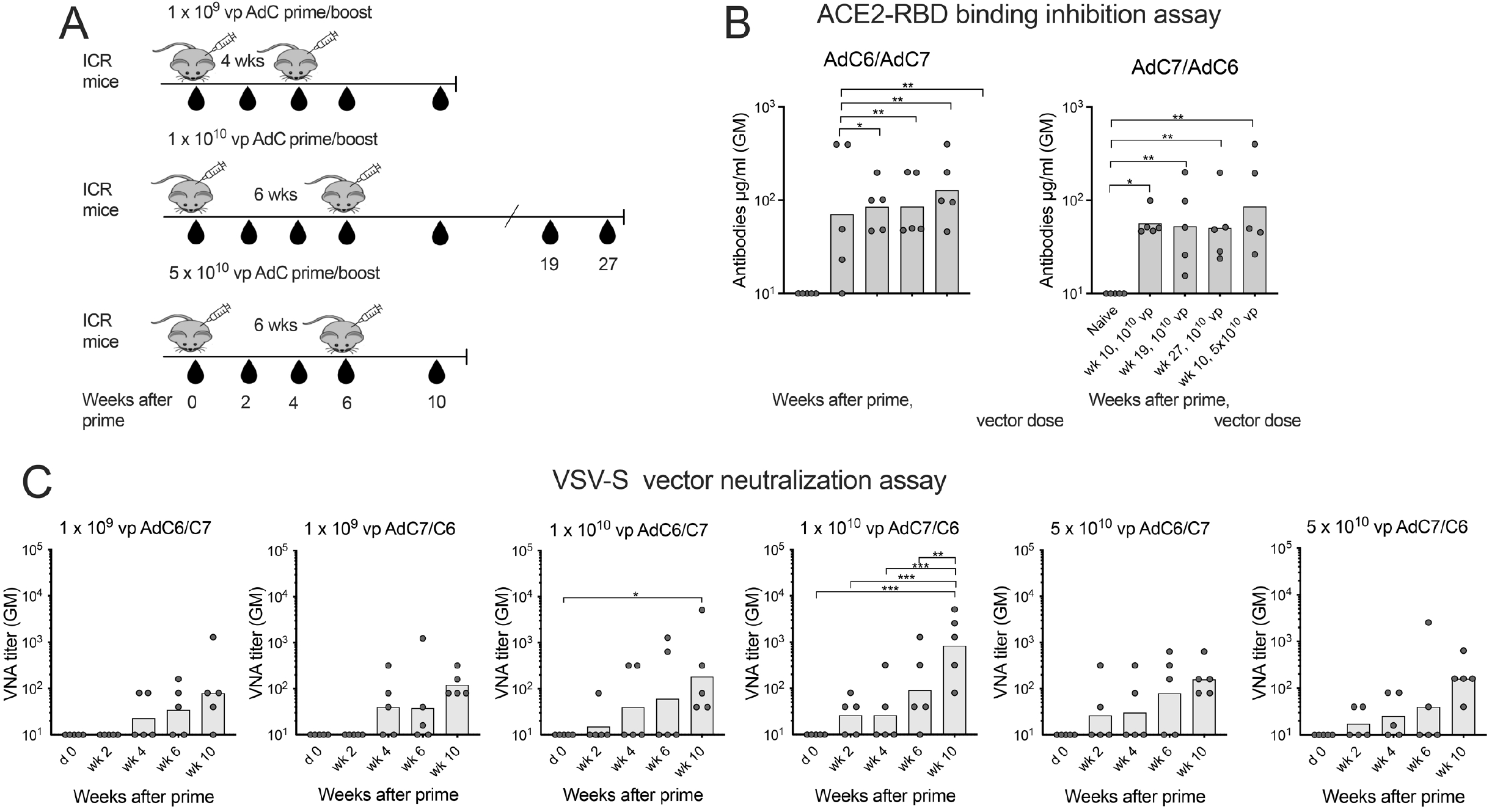
VNA responses to different dose of the AdC-S vectors. [A] Experimental outline. [B] Titers of antibody that inhibit ACE2 binding to the RBD domain of the S protein are shown for individual mice as μg of antibodies/ml calculated based on a standard. Negative data were set at 10 All samples tested after immunization had significantly higher levels of antibodies compared to sera from naïve mice as determined by multiple Mann-Whitney tests. [C] VNA titers to SARS-CoV-2 were tested with a S_SWE_ pseudotyped VSV vector. Titers are shown as circles for each mouse; bars show GMs. Negative data were set at 10. represent the serum dilution that achieved ≥ 50% inhibition of infection. Sera were tested starting at a 1:40 dilution. Significant differences were calculated by 2-way ANOVA with Tukey’s multiple comparison test.

Sera were in addition tested in a neutralization assay with the VSV-S_SWE_ vector. After the boost all mice but for one in the low dose AdC6-S_SWE_ group developed VNAs with geometric mean (GM) titers ranging from 1:80 in the low dose AdC6-S_SWE_ prime group to 1:840 in the intermediate dose AdC7-S_SWE_ prime group (Fig. 2C,D).

### Responses to different AdC-S vaccine regimens

We used different intervals between the prime and boost and explored the sequential use of homologous vs. heterologous AdC-S_SWE_ vectors. For these experiments 3 groups of 5 ICR mice were tested (Fig. 3A). The 1^st^ group was primed with 2 × 10^10^ vp of AdC6-S _SWE_. Mice were boosted 2 weeks later with the same dose of AdC7-S_SWE_. The 2^nd^ group received the same vaccines at the same dose and in the same order but the boost was delayed till week 8. The 3^rd^ group was primed with 2 × 10^10^ vp of AdC7-S_SWE_ and then boosted 8 weeks later with the same AdC7-S_SWE_ vector used at the same dose. Mice were bled periodically and antibody titers to S protein were determined by ELISA. As shown in Fig. 3B the early boost and the boost with the the homologous AdC vector only resulted in transient increases in S protein binding antibodies (Figure 1B). Even more pronounced differences were obtained with the neutralization assay (Fig. 3C), which showed that the early boost increased antibody titers 8-fold by week 10 (8 weeks after the boost), which then rapidly contracted. The homologous boost only achieved a 4-fold transient increase in VNA titers while the heterologous boost given 8 weeks after priming resulted by week 10 in an ~8-fold increase and then by week 15 in a 26-fold increase. The less potent recall response of the homologous prime boost was most likely caused by Ad-specific VNAs that developed after the prime against the homologous but not the heterologous AdC vector and which had reached a GM titer of 870 at the time of the homologous boost (data not shown).

**Fig. 3.**
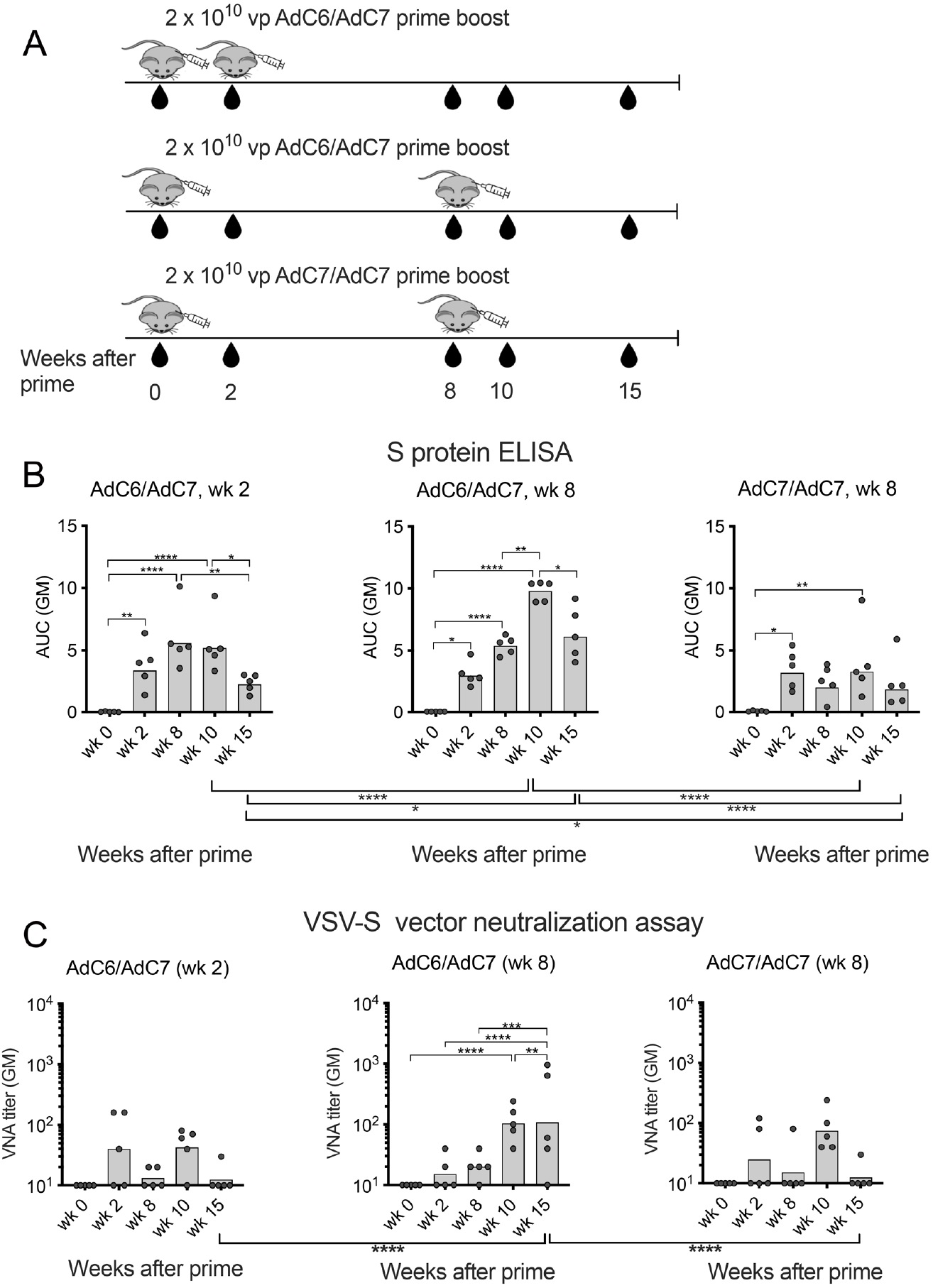
Factors that influence the effectiveness of a boost. [A] Experimental outline. [B] Antibody titers against S protein tested by ELISA. The design of the graphs mimics those of Figure 1C. Significant differences within one group are shown by lines and stars above each bar graph. Differences between the groups are shown below the x-axis. Both were calculated by 2-way ANOVA with Tukey’s correction. [C] VNA titers determined with a S_SWE_ pseudotyped VSV vector. Negative responses were set at 10. Significant differences were calculated by 2-way ANOVA with Tukey’s correction. Significant differences between samples within one group are shown above the bar graphs; those between groups are shown below the x-axis.

### Responses in young and aged mice

We conducted an experiment in young (6-8 weeks of age, n = 5) and aged (1-2 years of age, n = 7) C57Bl/6 mice. Mice were immunized with a low dose of 2 × 10^9^ vp of the AdC7-S_SWE_ vector and boosted 6 weeks later with 2 × 10^9^ vp of the AdC6-S_SWE_ vector (Fig. 4A). Naive young (n = 3) and aged (n = 2) mice were used as controls. Sera of individual mice were tested 2 weeks after the boost by an ELISA (Fig. 4B) and a VSV-S_SWE_ virus neutralization assay (Fig. 4C). Antibody responses tested by the S protein ELISA or the VNA assay tended to be higher in young mice although this did not to reach significance. All of the young mice developed VNAs while 2 out of 7 aged mice failed to respond.

**Fig. 4.**
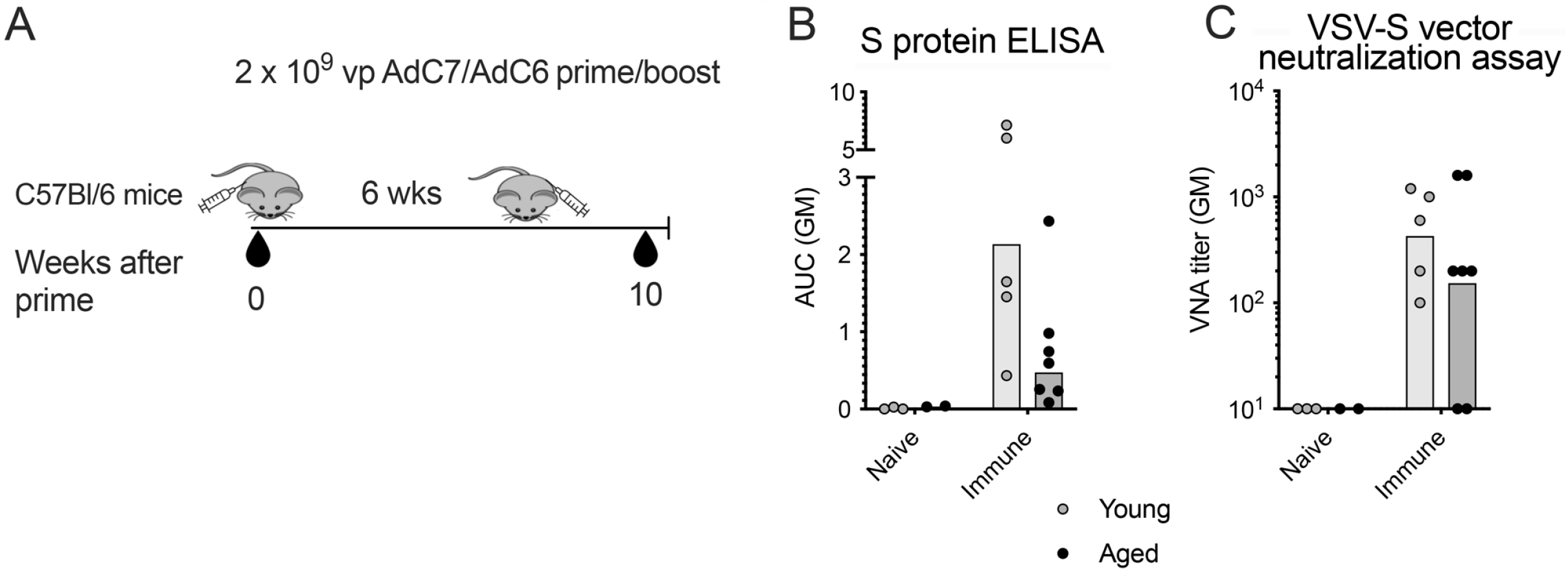
Antibody responses in young and old C57BI/6 mice. [A] Experimental outline. [B] Antibody titers at 4 weeks after the boost (wk 10) by ELISA shown as in Fig. 1C. VNA titers determined by a S_SWE_ pseudotyped VSV vector are shown as in Fig. 2C. There were no significant differences between the groups by multiple Mann-Whitney test.

### Cross-reactivity of vaccine-induced VNAs to SARS-CoV-2 variants

To test if the AdC-S vaccines induce antibodies that cross-neutralize viral variants we selected sera from ICR mice that had been immunized once or twice with AdC-S_SWE_ vectors and had shown positive neutralizition of the VSV-S_SWE_ vectors, and tested them for cross-reactivity with VSV vectors pseudotyped with different versions of S protein carrying RBD mutations found in viral variants of concern. A total of 61 or 22 sera harvested after priming (Fig. 5A) or boosting (Fig. 5B), respectively, were tested for cross-reactivity against VSV-S_SWE/B1.351_ and VSV-S_SWE/B1.1.7_ while 18 or 10 sera, respectively, were tested against VSV-S_SWE/B.1.617.2_. After priming, VNA titers were lower against VSV-S_SWE/B.1.1.7_ and VSV-S_SWE/B.1.6.17.2_ than against VSV-S_SWE_ and the latter difference reached significance. After the boost VNA titers became comparable. Sera with higher titers were, as expected, more cross-reactive than those with low titers. Accordingly, after priming antibody titers to S_SWE_ showed significant positive correlations to those against S_SWE/B.1.1.7_ and S_SWE/B.1.6.17.2_ and after the boost significance extended to correlations between S_SWE_ and S_SWE/B1.351_. A direct comparison of titers may not be valid as S protein-pseudotyped VSV vectors may exhibit differences in their sensitivity to antibody-mediated neutralization that could relate to factors other than the S protein sequences. We therefore also compared rates of responsiveness. All sera neutralize VSV-S_SWE_. Response rates against the variants were markedly lower after a single immunization: 20%, 40%, and 60% of sera failed to neutralize VSV-S_SWE/B.351_, VSV-S_SWE/B.1.1.7_, or VSV-S_SWE/B.1.617.2_, respectively. Reactivity markedly increased after the boost when ~80% of sera showed broad cross-reactivity (Fig. 5C,D).

**Fig. 5.**
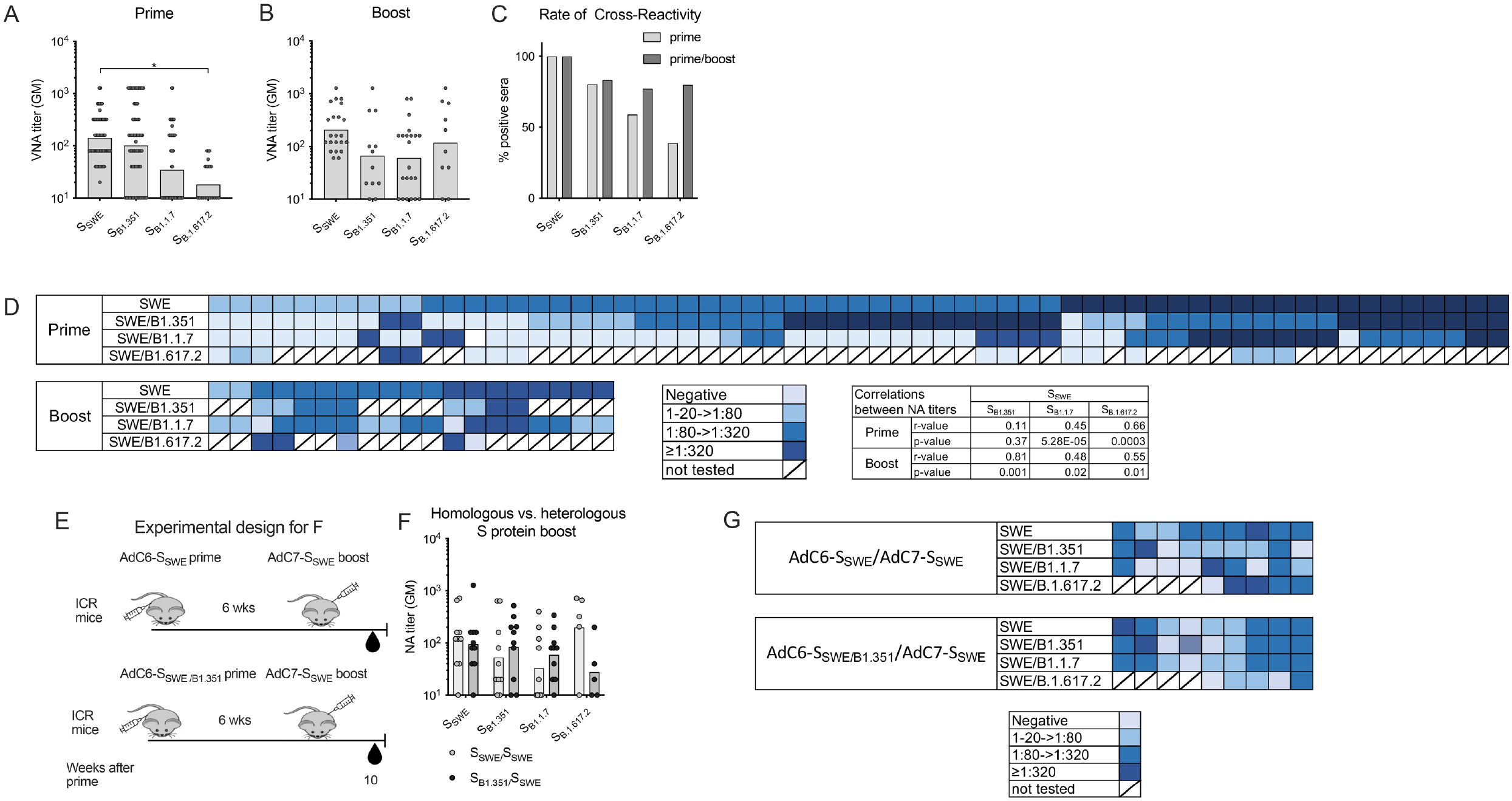
Cross-reactivity of AdC-S_SWE_ induced VNAs. Sera harvested after priming [A] or boosting [B] of ICR mice that had initially scored positive for neutralization of the VSV-S_SWE_ vectors were retested in parallel on this vector and on VSV-S_B1.351_, VSV-S_B1.17_ and VSV-S_B.1617.2_ vectors. [A, B] VNA titers in individual mice with bars indicating GMs. Negative responses were set at 10. Differences were calculated by one-way ANOVA [C] Percentages of sera that neutralized the different variants. [D] Heatmaps show levels of neutralization of the variants by individual sera. [E] Spearman correlations between titers against the immunizing variant VSV-S_SWE_ and the other variants. [E] Experimental design for the results shown in F and G. [F] Antibody titers that neutralized the different VSV-S vectors in individual mice tested 2 weeks after boosting. Boxes indicate GMs. There were no significant differences between the groups as determined by multiple Wilcoxon tests. [F] Heatmap shows levels of neutralization of the variants by individual sera.

To assess if we could increase cross-reactivity by using a prime boost regimen with vectors expressing two different S proteins we prime 10 mice either with 10^10^ vp of AdC6-S_SWE_ or AdC6-S_SWE/B.351_ vectors. Mice were boosted 6 weeks later with 10^10^ vp of AdC7-S_SWE_ vector and sera were tested 2 weeks later for neutralization of the different VSV-S vectors. Although there was a trend for higher VNA titers towards the VSV-S_SWE/B.1.1.7_ and VSV-S_SWE/B.1.6.17.2_ vectors upon sequential immunization with Ad vectors expressing two different S proteins, differences were subtle and failed to reach significance.

## DISCUSSION

In a remarkable feat of scientific ingenuity, vaccines were developed within weeks after SARS-CoV-2 was identified as the etiological agent of COVID-19. This was facilitated by earlier outbreaks with related viruses such as SARS-CoV-1 in 2002-2004^29,30^ or Middle East respiratory syndrome-related coronavirus (MERS-CoV), which appeared in 2012 in Saudi Arabia and has been circulating at low levels ever since.^31^ Experimental vaccines had been developed and tested pre-clinically against these coronaviruses^32–34^ and provided blueprints for the COVID-19 vaccines based on mRNAs or Ad vectors.

Protective immunity induced by mRNA vaccination against COVID-19 is short lived.^16–21,35^ As this was also observed in a previous trial testing an mRNA vaccine against rabies virus^36^ the rapid waning of protective immunity most likely relates to the vaccine platform rather than the disease application. The short-lived duration of mRNA vaccine-induced protection contrasts with that achieved by other viral vaccines based on attenuated viruses such as the those against smallpox or yellow fever virus, subunit vaccines such as the hepatitis B virus vaccine, or virus like particle vaccines such as the HPV vaccines, which in general provide protection for at least 5-10 years.^37–39^ The lifespan of an antibody secreting plasma cell and the magnitude of the memory B cell response is dictated by several factors including strength of B cell receptor signaling, which in part is driven by receptor cross-linking, presence of T help and the type of the inflammatory milieu that is elicited by a pathogen or a vaccine.^40^ COVID-19 mRNA vaccines were shown to result in germinal center-driven maturation of B cells^41^ but nevertheless induce mainly short-lived rather than long-lived plasma cells.^42^ In animal models Ad vectors were shown to induce germinal center B cells^43^ and very sustained plasma and memory B cell responses.^44^ Preliminary data indicate that this may translates into more sustained protection against COVID-19.^45,46^ mRNA vaccines have other disadvantages over Ad vector vaccines. mRNA vaccines are in general safe but rare serious adverse events in form of anaphylactic reactions or myocarditis have been reported.^47,48^ They are heat labile and must be stored at around −80°C and they are costly. Ad vector vaccines are heat stable and can be stored at 4°C.^49^ They are relatively inexpensive, which further facilitates their use in low-income countries. In general, Ad vectors are well tolerated although rare cases of potentially fatal cerebral venous sinus thrombosis (CVST) with thrombocytopenia have been linked to the J&J and AstraZeneca COVID-19 vaccines but not to the Sputnik V vaccine.^50,51^

One clear disadvantage of Ad vector vaccines is that immune responses to the transgene product are dampened by pre-existing VNAs to the vaccine carrier.^52–54^ Such antibodies can be induced by natural infections and consequently prevalence rates of VNAs to HAdV5 are globally high while VNAs to HAdV26 are common in humans residing in Africa.^52^ A single immunization with an Ad vector vaccine likely induces more modest levels of Ad-specific VNAs due to lower antigenic loads although repeated vaccination with the same Ad vectors is expected to gradually increase titers so that they will reach those seen after a natural infection. One clinical study with an HAdV5-based COVID vaccine from CanSino Biologicals monitored induction of Ad vector-specific VNAs and, as expected, showed that they increased after vaccination. The same study reported an albeit insignificant inverse correlation between VNA titers to the HAdV5 vector and COVID S protein-specific antibody responses.^55^ VNAs to the vaccine vectors will likely pose limitations to repeated booster immunization using the same Ad vector by not only reducing, but also, as we reported previously, modifying transgene product-specific immune responses.^56^

To avoid interference by Ad vector-specific VNAs, we developed a vaccine regimen composed of two serologically distinct E1-deleted AdC vectors. We selected AdC vectors as pre-existing VNAs to these chimpanzee viruses are rare in humans and individuals who have VNAs in general have low titers.^44^ We developed two serologically distinct AdC vectors to prevent blunting of booster immunizations by VNAs induced by the AdC vector used for priming. The two AdC vectors express the S protein of an isolate from Sweden, which is identical to that of the original Wuhan virus. We did not codon-optimize the S sequence, nor did we incorporate the K986P and V987P stabilizing mutations into the S2 sequence which both were used for the S gene expressed by the RNA vaccines,^57^ mutations of the S1/S2 furin cleavage site which, in addition to the stabilizing mutations, were incorporated into the S gene carried by the J&J COVID-19 vaccine,^58^ or AstraZeneca’s engineered leader sequence.^59^ Phase III trial results of the different Ad vector vaccines fail to indicate that any of these modifications have a major impact on vaccine efficacy.

In our study, as expected, the magnitude of S protein-specific antibody responses after a single immunization increased with higher vaccine doses. This pattern changed after boosting, where the highest dose performed poorly. Timing of boosting seems to affect the magnitude of the recall response as was reported for both the AstraZeneca and J&J vaccines.^60,61^ There were no significant differences between responses induced by the AdC6-S or AdC7-S vectors although after two immunizations the AdC7/AdC6 regimen tended to outperform the AdC6/AdC7 regimen. In our study a short interval of 2 weeks between two vaccine doses resulted only in marginal increases in antibody titers as compared to a longer interval of 8 weeks. We assume, but this must be investigated in more depth, that the optimal interval between repeated immunizations may depend on the vector dose used for priming with lower doses allowing for shorter intervals and may well exceed 8 weeks for higher doses.^62^ Repeated use of the same vector also reduces the effectiveness of the booster immunization, which is likely caused by Ad vector specific VNAs, which reduce transduction rates and thereby expression levels of the transgene product. Again, we assume that the inhibitory effect of Ad vector-specific VNAs will depend on pre-existing memory and vector dose, which both affect the magnitude of the response, as well as on timing of the booster immunization with longer intervals being advantageous by allowing for a decline in Ad vector-specific antibody titers. Alternatively, inhibition by Ad vector specific VNAs can be circumvented by using heterologous vectors for prime boosting such as in the Sputnik V vaccine or our pre-clinical vaccines or by changing vaccine platforms. In our study the importance of a boost was stressed by the relative lack of cross-reactivity of antibodies generated after a single immunization against S proteins with RBD mutations present in key variants such as the delta variant.

It is unlikely that SARS-CoV-2 will ever be eradicated. This will necessitate periodic booster immunizations and additional investments into 2^nd^ generation vaccines that achieve more sustained immunity and preferentially not only protection against symptomatic disease but also against infections. Booster immunizations with available Ad vector vaccine will over time become less and less effective due to the induction of vector neutralizing antibodies. This could be addressed using alterative Ad serotypes such as those described here.

## CONCLUSION

Two serologically distinct AdC vectors expressing S of an early SARS-CoV-2 isolate induce robust and sustained antibody responses upon prime-boosting that cross-neutralize viral variants. Additional work is needed test different doses and time intervals between prime boosting to further optimize this vaccine regimen.

## MATERIAL AND METHODS

### Cell lines

HEK 293 cells and VeroE6 cells were grown in Dulbecco’s Modified Eagles medium (DMEM) supplemented with 10% fetal bovine serum (FBS) and antibiotics in a 5% CO_2_ incubator. BHK-21/WI-2 cells were grown in DMEM supplemented with 5% FBS and antibiotics at 5% CO2.

### Expression of ACE-2

Cells diluted to 1 × 10^6^ per microtiter plate well were incubated for 45 min at room temperature with a 1 in 100 dilution of a mouse IgG2a antibody to human/hamster ACE-2 (clone 171606 R&D Systems, Minneapolis, MN). After washing, cells were incubated for 45 min with an FITC-labeled goat-anti mouse IgG and a live cell stain. After washing, cells were analyzed by flow cytometry.

### Generation of and quality control AdC-S vectors

The cDNA sequence for S _SWE_ of SARS-CoV-2 was obtained from the laboratory of Dr. Elledge, Massachusetts General Hospital, Boston, MA (GenBank: QIC53204.1). Sequencing revealed a frame-shift mutation that was corrected by site-directed mutagenesis. The corrected S gene was cloned into a shuttle vector and from there into the viral molecular clones of AdC6 and AdC7. Viral vectors were rescued, expanded, and purified on HEK 293 cells as described.^26^ Viral DNA was isolated and tested by restriction enzyme digest for presence and integrity of the inserts. Viral titers were determined by spectrophotometry. Yields for AdC6-S and AdC7-S were 1.4 × 10^13^ and 2.5 × 10^13^ respectively upon expansion in 2 × 10^8^ HEK293 cell. Genetic stability was determined by restriction enzyme digest of purified viral DNA after 12 sequential passages of the vectors.

To allow for rapid exchange of the RBD sequence we inserted an AgeI site into position 1640 of the S gene which changes the nucleotide but not the amino acid sequence of S. The variant RDB sequences including flanking regions and convenient restriction enzyme sites were synthesized (Genscript Biotech, Piscataway, NJ) and cloned into the S_SWE_ gene using restriction enzymes BsrGI which cuts at base pair 1104 and Age1 of the S_B1.351_ sequence and BseGI and NheI, which cuts at base pair 2015 of the S_B1.1.7_ and S_B.1.617.2_ sequences.

### Generation of VSV-S vectors

VSV pseudotyped with S of SARS-CoV-2 were generated in BHK-21/WI-2 cells using a the ΔG-GFP (G*ΔG-GFP) rVSV kit (Kerafast, Boston, MA, USA) and S sequences cloned into an expression plasmid under the control of the CMV promoter as described previously.^66^ BHK-21/WI-2 cells were plated into a 6 well plate, the next day cells were first transfected with 1 μg/well of the paT7 plasmids using polyethyleneimine (PEI) (Polysciences, Inc. Warrington, PA); 2 hours later, the cells were treated with a predetermined optimal amount of ΔG-GFP (G*ΔG-GFP) rVSV. After a 24-hr incubation at 37°C in a 5% CO_2_ incubator, supernatants were harvested, aliquoted, and stored at −80 °C.

### Protein expression

The AdC-S vectors were tested for protein expression upon transfection of HEK 293 cells with 1000 vp of the vectors for 48 hr at 37°C. Cells were lysed in RIPA buffer supplemented with a 1%μl protease inhibitor (Santa Cruz Biotechnology Inc., Dallas, TX). A 15 ul of lysate was resolved on 12% SDS-PAGE and transferred to a polyvinylidene difluoride (PVDF) membrane (Merck Millipore, Burlington, MA). The membrane was blocked in 5% powder milk overnight in 4°C. The primary antibody to S diluted to 1:1000 in saline (clone ABM19C9, Abenomics, San Diego, CA) was added for 1 hr at room temperature. Membranes were washed with 1X TBS-T prior to incubating with HRP-conjugated goat anti-rabbit secondary IgG (ab6721, Abcam, Cambridge UK) for 1 hr at room temperature. In parallel the membrane was probed with a mouse monoclonal IgG antibody to ß-actin (Sc-47778, Santa Cruz Biotechnology, Dallas, TX) as a loading control for 1 hr at room temperature. The loading control antibody was probed with HRP-conjugated goat anti-mouse secondary IgG (SAB3701047, Sigma, St. Louis, MO) for 1 hr at room temperature. Membranes were washed 3 times with 1X TBS-T. The developing agent Super Signal West Pico Chemiluminescent (Thermo Fisher Scientific, Waltham, MA) was added. Membranes were shaken in the dark for 5 min, dried and developed.

### Mice and mouse procedures

Female C57Bl/6 mice (6–8 weeks of age) were purchased from the Jackson Laboratories (Bar Harbor, ME). Outbred 6–8-week-old female ICR mice were obtained from Taconic Biosciences (New York, NY). Mice were housed at the Wistar Institute Animal Facility. All mouse procedures followed approved protocols. Mice were vaccinated intramuscularly (i.m.) with the AdC vectors diluted in 200 μl of sterile saline. Mice were bled from the saphenous *vein* and blood was collected into 4% sodium carbonate and Liebowitz’s-15 (L-15) medium. Serum was isolated 30 min later upon a 10 min centrifugation of tubes at 14000 rpm.

### ELISA

Sera from individual mice were tested for S-specific antibodies by ELISA on plates coated overnight with 100 μl of a mixture of S1 and S2 (Native Antigen Company, Kidlington, UK) each diluted to 1 μg/ml in bicarbonate buffer. The next day plates were washed and blocked for 24 hr at 4°C with 150 μl of a 3% BSA-PBS solution. Sera were diluted in 3%BSA-PBS and 80μl of the dilutions were added to the wells after the blocking solution had been discarded. Plates were incubated at room temperature for 1 hr and then washed 4 x with 150 μl of PBS. An alkaline-phosphate-conjugated goat anti-mouse IgG antibody (Sigma-Aldrich, St Louis, MO) diluted to 1:1500 in 3%BAS-PBS was added at 60 μl/well for 1 hr at room temperature. Plates were washed 4 times and substrate composed of phosphatase substrate tables (Sigma-Aldrich, St Louis, MO) diluted in 5 ml of diethanolamine buffer per tablet was added. Plates were read in an ELISA reader at 405 nm. Coated wells only receiving the substrate served to determine background. Background data were subtracted from the experimental data.

### VNA assay with VSV-S

VSV-S vectors was initially titrated on BHK-21/WI-2 cells. The cells were diluted to 4 × 10^5^ cells/ml in DMEM with 5%FBS and 5 μl were of the cell dilution was added to each wells of Terasaki plates. The next day serial dilutions of VSV-S vectors were added to duplicate wells. The following day numbers of fluorescent cells/well were counted with a fluorescent microscope (Suppl. Fig. 1D). For the neutralization assay a dose of VSV that infected ~40-60 cells/well was selected.

For the neutralization assay, BHK-21/WI-2 cells were plated into Terasaki plate wells as described above. The following day, sera were serially diluted in DMEM with 5% FBS and then incubated with VSV-S diluted in the same medium for 90 min at room temperature under gentle agitation starting with a serum dilution of 1 in 40 or 1 in 100. 5μl aliquots of the mixtures were transferred onto BHK-21/WI-2 within the Terasaki plate wells. Each serum dilution was tested in duplicates, an additional 4-6 wells were treated with VSV-S that had been incubated with medium rather than serum. Plates were incubated for 24-48 hr and then numbers of fluorescent cells were counted. Titers were set as the last serum dilution reduced numbers of green-fluorescent cells by at least 50%.

The COV-PosSet-S from RayBiotech (Peachtree Corners, GA), which contains 20 human samples from individuals that recovered from COVID-19 or from uninfected individuals was used to validate the neutralization assay.

### ACE binding inhibition assay

Sera were tested for inhibition of ACE binding to the RBD of S by the Anti-SARS-CoV-2 Neutralizing Antibody Titer Serologic Assay Kit from Acro Biosystems (Newark, DE) following the manufacturer’s instructions. A positive standard with a known concentration provided by the kit was used to extrapolated anti-S antibody concentrations in μg in the mouse sera.

## ACKNOWLEDGEMENTS

This work was funded by grants from The G. Harold and Leila Y. Mathers Charitable Foundation, the Commonwealth of Pennsylvania, and the Wistar Science Discovery Fund. MH was the recipient from the Fellowship from Janssen Scientific Affairs. Support for Shared Resources utilized in this study was provided by Cancer Center Support Grant (CCSG) P30CA010815 to The Wistar Institute.

---

**Supplemental Fig. 1.**
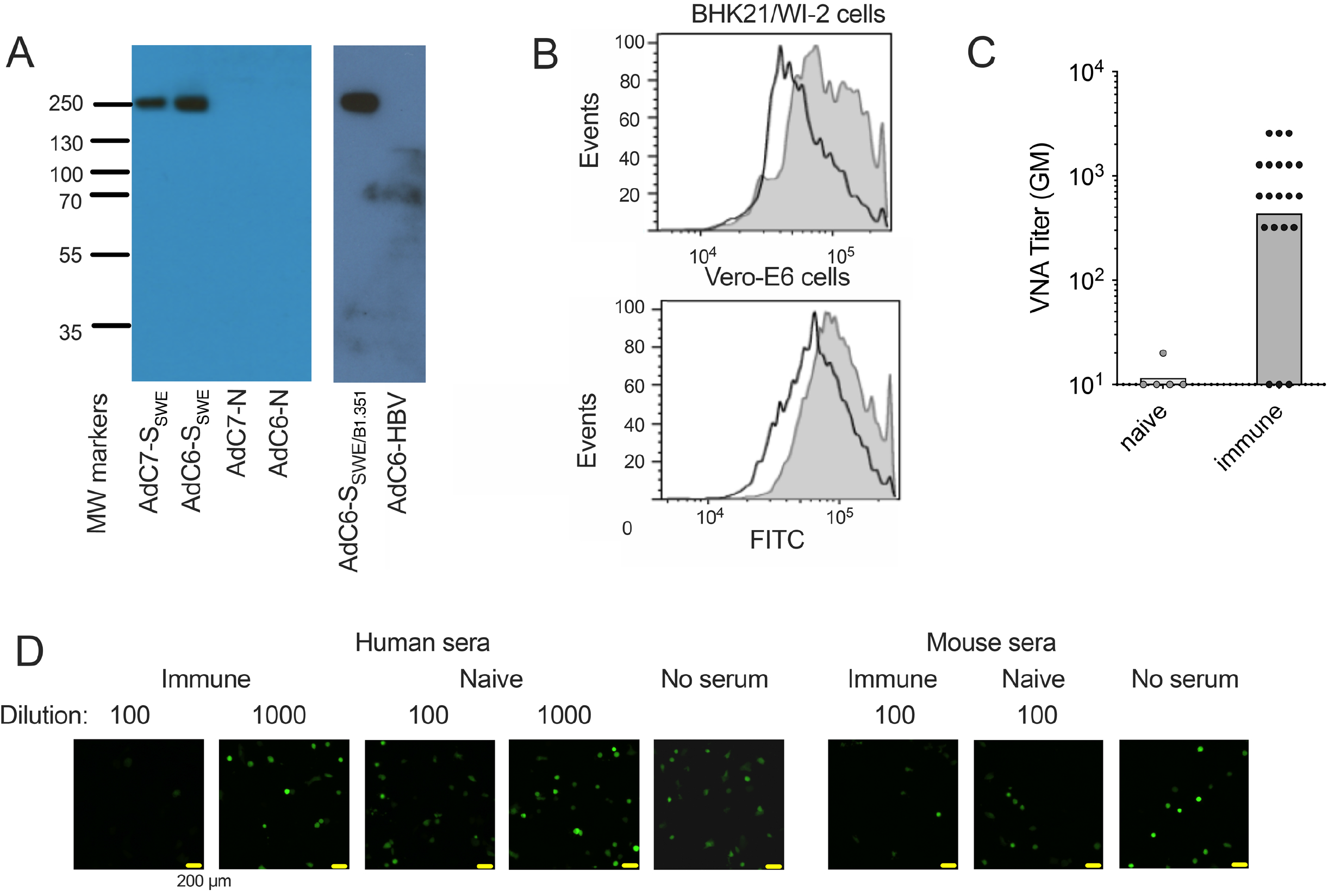
[A] Expression of the S protein by the AdC6-S_SWE_, AdC7-S_SWE_ and AdC6-S_SWE/B1.351_ vectors tested by Western blots of cells infected with the vectors or Ad vectors expressing an unrelated transgene product (N or HBV) as controls. MW - molecular weight marker. [B] Flow blots of the indicated cells after they were stained with an FITC-labeled antibody to ACE2 (filled grey) or an isotype control (bold black line) antibody. [C] Results of a neutralization assay using a panel of commercially available human SARS-CoV-2 immune or naïve sera and the VSV_SWE_ vector on BHK21/WI-2 cells. [D] Fluorescent staining of VSV-S_SWE_ transduced BHK21/WI-2 cells. Immune and naïve human sera were tested at 1:100 and 1:1000 dilutions while mouse sera were tested at a 1:100 dilution. Wells that received VSV-S_SWE_ served as controls.

